# Spatial transcriptomic analysis of virtual prostate biopsy reveals confounding effect of heterogeneity on genomic signature scoring

**DOI:** 10.1101/2023.03.08.531491

**Authors:** Sandy Figiel, Wencheng Yin, Dimitrios Doultsinos, Andrew Erickson, Ninu Poulose, Reema Singh, Anette Magnussen, Thineskrishna Anbarasan, Renuka Teague, Mengxiao He, Joakim Lundeberg, Massimo Loda, Clare Verrill, Richard Colling, Pelvender S. Gill, Richard J. Bryant, Freddie C. Hamdy, Dan J. Woodcock, Ian G. Mills, Olivier Cussenot, Alastair D. Lamb

## Abstract

Genetic signatures have added a molecular dimension to prognostics and therapeutic decision-making. However, tumour heterogeneity in prostate cancer and current sampling methods could confound accurate assessment. Based on previously published spatial transcriptomic data from multifocal prostate cancer, we created virtual biopsy models that mimic conventional biopsy placement and core size. We then analysed the gene expression of different prognostic signatures (OncotypeDx^®^, Decipher^®^, Prostadiag^®^) using a step-wise approach increasing resolution from pseudo-bulk analysis of the whole biopsy, to differentiation by tissue subtype (benign, stroma, tumour), followed by distinct tumour grade and finally clonal resolution. The gene expression profile of virtual tumour biopsies revealed clear differences between grade groups and tumour clones, compared to a benign control, which were not reflected in bulk analyses. This suggests that bulk analyses of whole biopsies or tumour-only areas, as used in clinical practice, may provide an inaccurate assessment of gene profiles. The type of tissue, the grade of the tumour and the clonal composition all influence the gene expression in a biopsy. Clinical decision making based on biopsy genomics should be made with caution while we await more precise targeting and cost-effective spatial analyses.

**Patient summary:** Prostate cancers are very variable, including within a single tumour. Current genetic scoring systems, which are sometimes used to make decisions for how to treat patients with prostate cancer, are based on sampling methods which do not reflect these variations. We found, using state-of-the-art spatial genetic technology to simulate accurate assessment of variation in biopsies, that the current approaches miss important details which could negatively impact clinical decisions.

**Take home message:** Virtual biopsies from spatial transcriptomic analysis of a whole prostate reveal that current genomic risk scores potentially deliver misleading results as they are based on bulk analysis of prostate biopsies and ignore tumour heterogeneity.

---

Prostate cancer (PCa) progression is unpredictable. Some tumours are indolent with a low propensity for progression, whilst others tumours are aggressive with rapid progression and metastasis [1]. Recent prognostic genetic signatures have added a molecular dimension to therapeutic decision-making [2,3]. However, a drawback of biopsy-derived molecular signatures is inaccurate sampling and failure to properly consider intra-tumoral heterogeneity [2,4]. In view of this, we compared three different prognostic signatures (OncotypeDx^®^[5], Decipher^®^[6] and ProstaDiag^®^[7]) in virtual biopsy models using spatial transcriptomics (ST). These signatures include genes associated with biological processes that may be fundamental to tumour development. In evaluating these signatures, we assessed gene expression assigned to these consensus processes.

Gene expression analysis was performed on a tissue sample obtained by radical prostatectomy from a patient with multifocal PCa. We used our recently published organ-wide ST data [8] to construct virtual biopsy models that mimic conventional biopsy placement and core size (**Fig. 1**). The biopsy regions were intentionally positioned to represent maximum heterogeneity. All spots containing less than 500 Unique Molecular Identifier counts were removed. Genes detectable at this threshold in less than 10% of the spots were also removed. Overall, we examined the expression of 9 of the 12 OncotypeDx^®^ genes, 11 of the 19 Decipher^®^ genes, and 28 of the 36 Prostadiag^®^ genes, excluding housekeeping genes. We extracted gene expression data from the barcode ID of designed biopsies (LoupeBrowser, 6.3.0) using R (4.22). In all analyses, we normalised the libraries using spaceranger aggr (2.0). Fold changes with false discovery rate were analysed using *EdgeR* (3.40.1).

**Figure 1:**
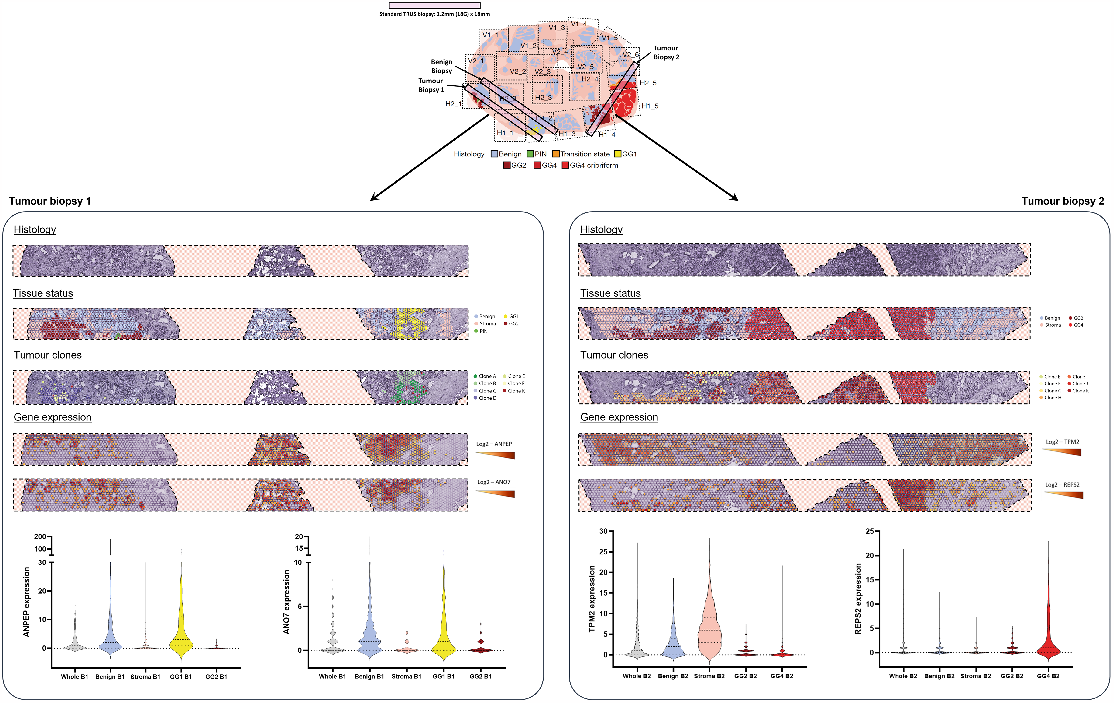
Spatial visualisation of virtual biopsies. We used our recently published organ-wide spatial transcriptomic data[8] to construct virtual biopsy models (2 tumour biopsies and 1 benign biopsy) that mimic conventional biopsy placement and core size. Visualisation of histology and tissue status (GG: Gleason grade group; PIN: prostatic intraepithelial neoplasia) and tumour clones from each tumour biopsy. Spatial visualisation of gene expression (ANPEP, ANO7, TPM2 and REPS2) in each tumour biopsy. Violin plots representing gene expression according to histological status. B1: tumour biopsy 1; B2: tumour biopsy 2.

We compared the gene expression of virtual tumour biopsies to a benign control using increasing resolution: from whole biopsy through tissue subtype (benign, stroma, tumour) and tumour grade to clonal level (**Fig. 1**). A consensus pathology underpins this analysis, with two pathologists independently annotating each 55µm ST spot (approximately 10-15 cells).

We observed clear evidence of variation in the expression of constituent genes from different prognostic signatures within biopsies (**Fig. 2**). In tumour biopsy 1, the expression of the cellular organisation markers (FLNC, TPM2, GSN) of OncotypeDx^®^ signature was decreased in a Gleason Grade Group (GG) 1 area (logFC=-0.91; -1.05; -0.68 respectively), while it was increased in the region of GG2 cancer (logFC=0.73; 1.09 respectively), compared to the control biopsy. Importantly, the distinction would be lost if either the whole biopsy, or tumour areas alone, were analysed in bulk (**Fig. 2A**). We observed similar results when we extended these analyses to other prognostic signatures (**Supplementary Fig. 1A & 2**). Further differences in gene expression were found between GG1 and GG2 cells compared to the control, such as the expression of genes involved in epithelial-mesenchymal transition (ANPEP, COL1A1, COL1A2, FMOD, SPARC), transport (ANO7, CHRNA2) and cell cycle (NCAPD3, ZWILCH).

**Figure 2:**
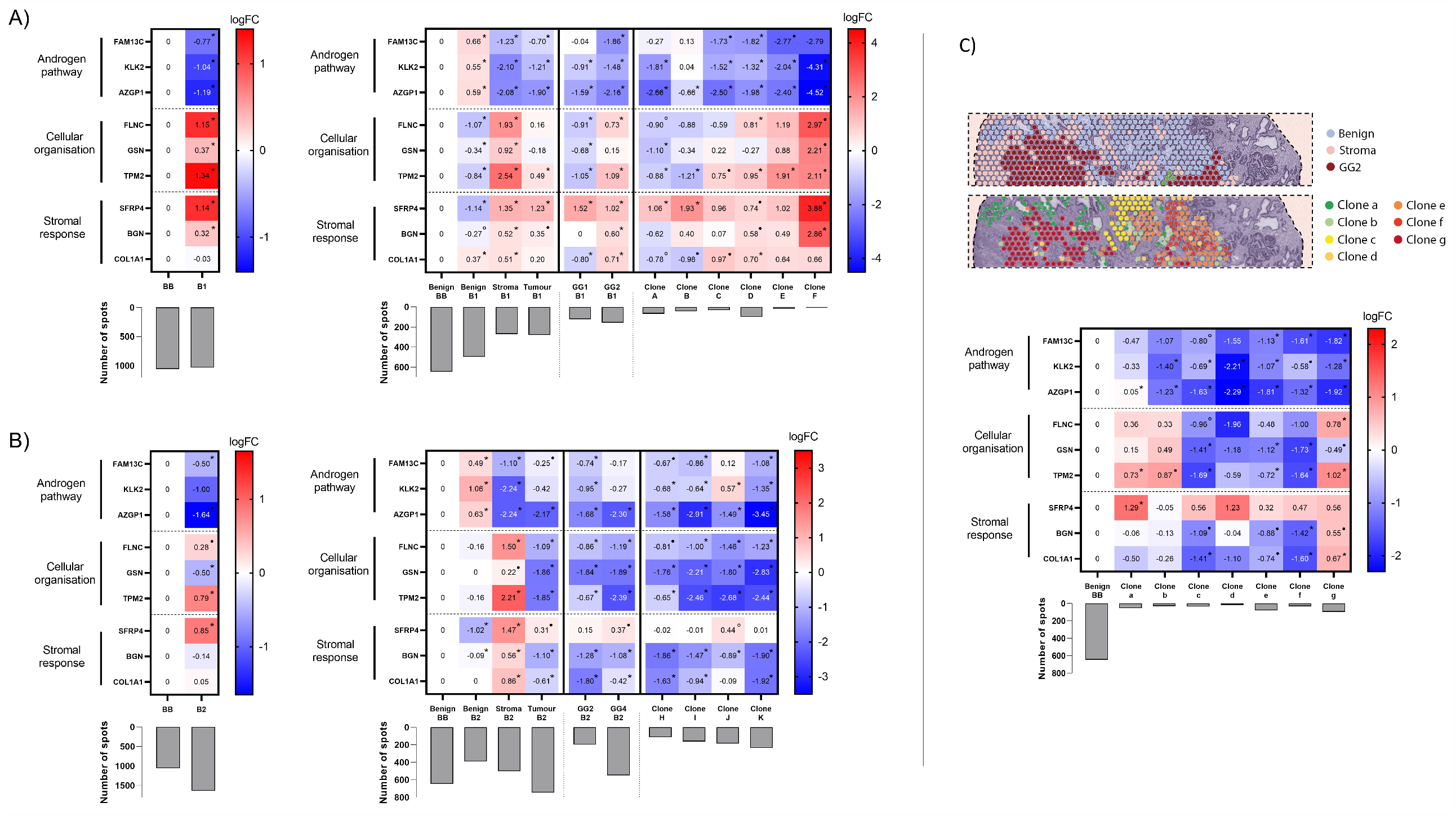
Gene expression profile of the OncotypeDx^®^ signature. Heatmap of gene expression (logFC) of the OncotypeDx^®^ signature at different levels of precision (whole biopsy, tissue subtype, tumour grade and clonal level) in tumour biopsy 1 (A), tumour biopsy 2 (B) and section H2_1 (C). The histograms represent the number of spatial transcriptomic spots for each entity. False discovery rate (FDR) is indicated: *FDR < 0.01; ^•^FDR < 0.05, °FDR < 0.1. FC: fold change; BB: benign biopsy; B1: tumour biopsy 1; B2: tumour biopsy 2; GG: Gleason grade group.

Previously [8], we interrogated the expression data using spatial inferred copy number variations and constructed a phylogenetic-tree to describe sequential clonal events in tumour regions of the tissue. In this work, we also performed clonal-level analyses based on the previously identified 11 clonal groups of tumour cells (clone A to clone K) [8]. In addition, we found expression differences between clones of the same histological grade, further supporting the need to respect heterogeneity in interpreting genomic scores (**Fig. 2A; Supplementary Fig. 1A & 2**).

In tumour biopsy 2, the expression of the cellular organisation markers (FLNC, TPM2) and the stromal response markers (BGN, COL1A1) decreased in GG2/GG4 areas compared to the benign control (logFC=-0.86/-1.14; -0.62/-2.39; -1.28/-1.08; -1.80/-0.42 respectively). Conversely, these genes demonstrated an inverse profile if the biopsy was analysed in bulk (logFC=0.28; 0.79; -0.14; 0.05) (**Fig. 2B**). Therefore, the gene expression profile of biopsy 2 as a whole, which comprises 46% tumour tissue and 31% stroma, did not reflect the gene expression profile of the tumour areas. The tumour profile was masked by the stroma profile. In addition, the expression of the androgen pathway markers (FAM13C, KLK2) of OncotypeDx^®^ were decreased in a region of GG2 cancer (logFC=-0.74; -0.95 respectively) but not in the GG4 area (logFC =-0.17; -0.27 respectively) (**Fig. 2B**). Similar to Biopsy 1, this information would be missed if either the whole biopsy, or tumour areas alone, were analysed in bulk. We obtained consistent results when we extended these analyses to the other genomic scores. As with biopsy 1, we also observed differences between tumour clones (**Supplementary Fig. 1B & 3**).

Given our previous finding of early cancer-associated events in histo-pathologically benign prostate tissue (section H2_1) [8], we proceeded to a detailed examination of this region (**Fig. 2C; Supplementary Fig. 4**). We interrogated the expression of the different signatures in the 7 clonal groups previously identified: clones a and c comprise 100% benign cells; clones d to f 25% and clone g less than 25%. The gene expression profile of clone c, previously identified as altered benign, was closer to that of tumour clones d, e and f than to that of benign clones a and b, in the 3 prognostic signatures studied. For the OncotypeDx^®^ signature, markers of cell organisation (FLNC, GSN, TPM2) and stromal response (BGN and COL1A1) decreased in clone c cells (logFC=-0.96; -1.41; -1.69; -1.09; -1.41 respectively), in clear distinction from clones a/b (logFC=0.36/0.33; 0.15/0.49; 0.73/0.87; -0.06/-0.13; -0.50/-0.26 respectively) compared to the benign control. These interesting findings from an area of altered benign tissue suggest an intermediate stage between benign and malignant cells, which would not have been identified in a bulk analysis.

Other studies have reported the co-founding effects of heterogeneity in genomic tests [2,9,10]. Despite their high cost, we believe that ST analyses provide a comprehensive *in situ* transcriptional assessment of heterogeneity.

We recognise the limitations of presenting data from a single patient. Furthermore, there are other examples of molecular heterogeneity (e.g. point mutations and epigenetic changes) that are not captured by the ST approach. The current cost for organ-scale analysis of a single prostate is around £100,000, but as such costs fall, and we put more prostates through our pipeline, we anticipate developing a more comprehensive understanding of PCa heterogeneity and lethal clonality.

In conclusion, we show that tissue type, tumour grade, and clonal composition of tissue all influence gene expression within a biopsy sample. Bulk analyses of prostate biopsies for genomic scores used in clinical practice effectively ignore these distinctions. To maximise the potential of biopsy-based genomics in clinical decision making, we believe that precise targeting will need to be combined with more granular spatial analyses, in order to provide accurate scores which will better-inform clinical decisions.

## Acknowledgements

This project has received funding from the European Research Council (ERC) under the European Union’s Horizon 2020 research and innovation programme (grant agreement no. 101021019). The study was also supported by the Swedish Cancer Society, Swedish Foundation for Strategic Research, AstraZeneca and Science for Life Laboratory. We also acknowledge the Swedish Childhood Tumour Biobank, supported by the Swedish Childhood Cancer Fund, for access and handling of patient biobank material/sequencing data and the Swedish Childhood Cancer Fund. We would like to thank the National Genomics Infrastructure, Sweden, for providing infrastructure support. We thank A. Mollbrink, X.M. Abalo, M. Nistér and P. Lundin for helpful assistance and discussions. A.D.L. was supported by a Cancer Research UK Clinician Scientist Fellowship award (C57899/A25812) that also funded W.Y, A.E., F.C.H, R.C. and A.D.L. have received support from the Oxford National Institute for Health Research (NIHR) Biomedical Research Centre Surgical Innovation and Evaluation Theme. S.F was funded by the Hanson Trust Research (HJD00160). I.G.M. is grateful to the John Black Charitable Foundation for support, D.D is grateful to the Prostate Cancer Foundation, and D.J.W. is grateful to the Cancer Research UK Oxford Centre. Computation used the Oxford Biomedical Research Computing facility, a joint development between the Wellcome Centre for Human Genetics and the Big Data Institute supported by Health Data Research UK and the NIHR Oxford Biomedical Research Centre. The views expressed are those of the author(s) and not necessarily those of the NHS, the NIHR or the Department of Health. We appreciate the kind assistance of A. Ståhl, R.A. Novoa and K. Rieger with additional histology assessment of specimen in this study.

## Competing interests

M.H. and J.L. are scientific consultants to 10x Genomics, Inc.

