## Supplementary Fig. 1 for "Spatial transcriptomic analysis of virtual prostate biopsy reveals confounding effect of heterogeneity on genomic signature scoring"

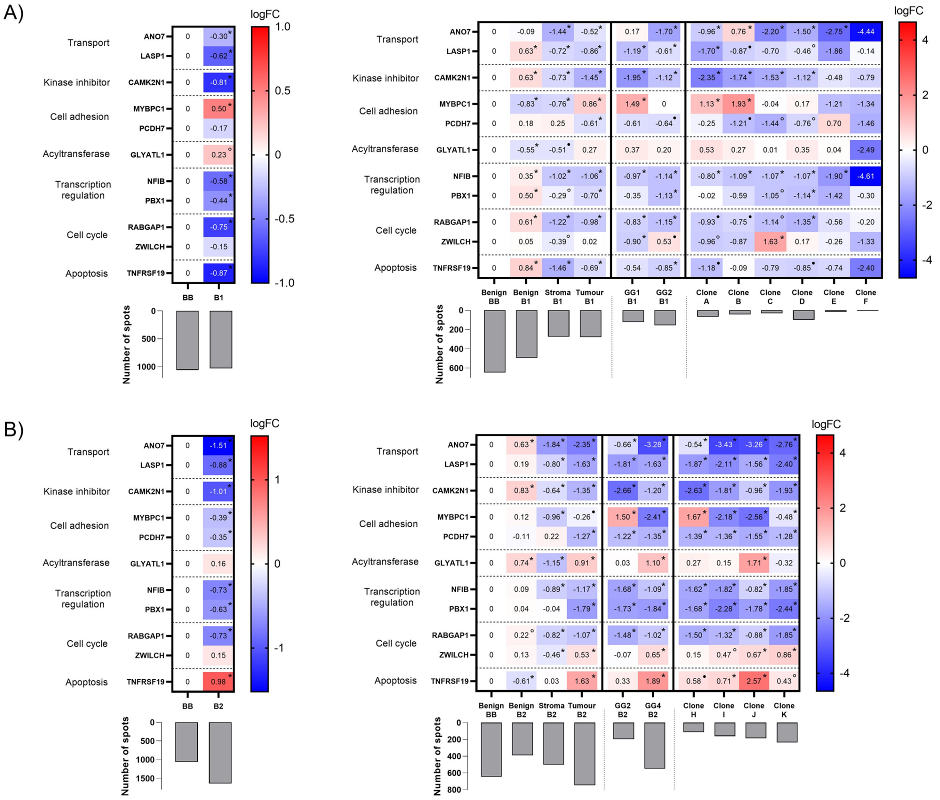


**eFigure 1: Gene expression profile of the Decipher® signature.**

Heatmap of gene expression (logFC) of the Decipher® signature at different levels of precision (whole biopsy, tissue subtype, tumour grade and clonal level) in tumour biopsy 1 (A) and tumour biopsy 2 (B). The histograms represent the number of spatial transcriptomic spots for each entity. False discovery rate (FDR) is indicated: *FDR < 0.01; •FDR < 0.05, °FDR < 0.1. FC: fold change; BB: benign biopsy; B1: tumour biopsy 1; B2: tumour biopsy 2; GG: Gleason grade group.
