## Supplementary Fig. 3 for "Spatial transcriptomic analysis of virtual prostate biopsy reveals confounding effect of heterogeneity on genomic signature scoring"

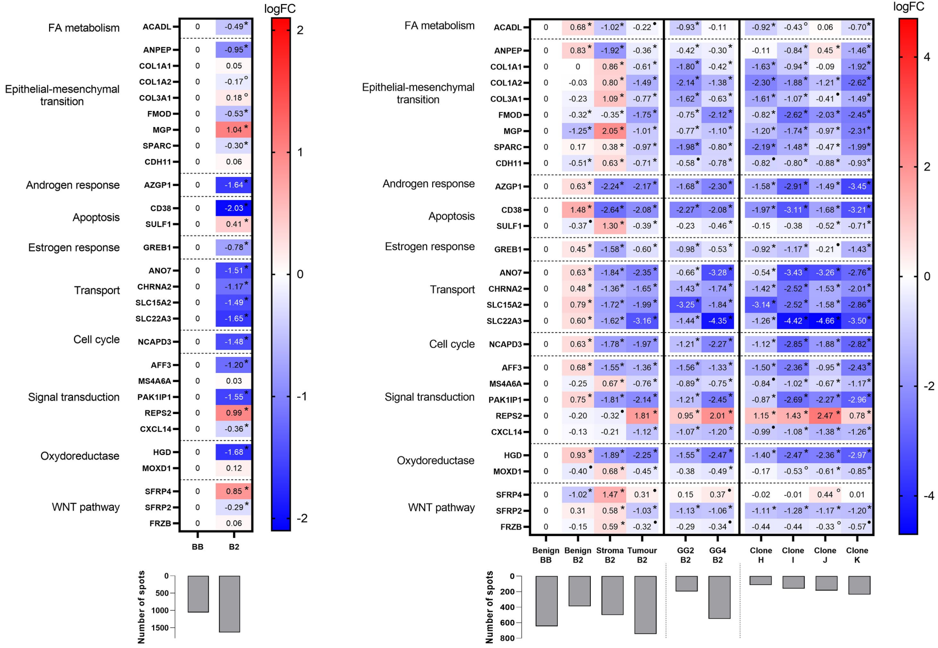


**eFigure 3: Gene expression profile of the Prostadiag® signature in tumour biopsy 2.**
Heatmap of gene expression (logFC) of the Prostadiag® signature at different levels of precision (whole biopsy, tissue subtype, tumour grade and clonal level) in tumour biopsy 2. The histograms represent the number of spatial transcriptomic spots for each entity. False discovery rate (FDR) is indicated: *FDR < 0.01; •FDR < 0.05, °FDR < 0.1. FC: fold change; BB: benign biopsy; B2: tumour biopsy 2; GG: Gleason grade group.
