## Supplementary Fig. 4 for "Spatial transcriptomic analysis of virtual prostate biopsy reveals confounding effect of heterogeneity on genomic signature scoring"

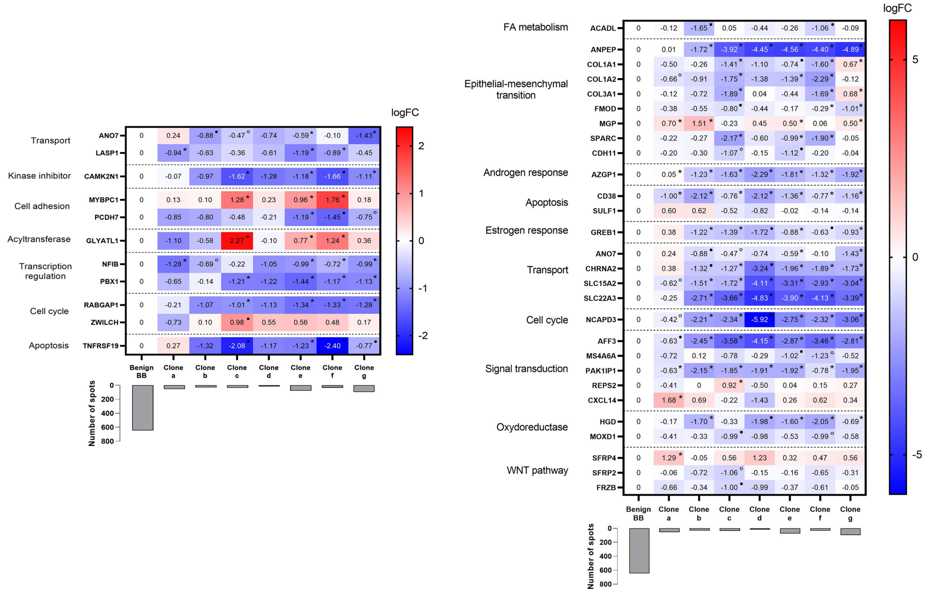


**eFigure 4: Gene expression profile of the Decipher® and Prostadiag® signature in section H2_1.**
Heatmap of gene expression (logFC) of the Decipher and Prostadiag® signature at the clonal level in section H2_1. The histograms represent the number of spatial transcriptomic spots for each entity. False discovery rate (FDR) is indicated: *FDR < 0.01; •FDR < 0.05, °FDR < 0.1. FC: fold change; BB: benign biopsy.
